# Inverse Agonism of the FFA4 free fatty acid receptor controls both adipogenesis and mature adipocyte function

**DOI:** 10.1101/2025.02.02.636098

**Authors:** WS Alshammari, EM Duncan, L Vita, M Kenawy, B Dibnah, M Wabitsch, G Gould, BD Hudson

**Affiliations:** Centre for Translational Pharmacology, School of Molecular Bioscience, University of Glasgow, Glasgow, UK; German Center for Child and Adolescent Health (DZKJ), Division of Pediatric Endocrinology and Diabetes, Department of Pediatrics and Adolescent Medicine, Ulm University Medical Center, Germany; Institute of Pharmacy and Biomedical Sciences, University of Strathclyde, Glasgow, UK

**Keywords:** Free fatty acid, Adipocyte, FFA4 receptor, adipogenesis, glucose uptake, lipolysis

## Abstract

Adipocyte disfunction is an important component of many metabolic disorders and there is a need for pharmacological approaches that can restore normal adipocyte function. The FFA4 receptor is a G protein coupled receptor (GPCR), activated by long chain free fatty acids (FFAs), that controls adipocyte function. Importantly, adipocytes produce FFAs, which may directly activate FFA4 and there is a need to better understand how FFAs produced by adipocytes interact with FFA4 signalling. In this study we have employed human and mouse adipocyte cell models to determine how pharmacological agonism or antagonism of FFA4 affects adipogenesis, lipolysis and glucose uptake. We show that a commonly used FFA4 antagonist, AH7614, is an inverse agonist and that treating adipocytes with this compound suppressed adipogenesis, inhibits glucose uptake and enhances isoprenaline stimulated lipolysis. In contrast, treatment with a synthetic FFA4 agonist, TUG-891, has only modest effects on adipogenesis and lipolysis, while showing no effect on glucose uptake. To explore the mechanism for why AH7614 but not TUG-891 affects adipocyte function, we demonstrate that during adipogenic differentiation sufficient FFAs are released into the culture medium to activate FFA4, suggesting AH7614 inhibits an autocrine feedback loop to suppress adipogenesis. In contrast, during lipolysis experiments, insufficient FFAs were released to activate the receptor, suggesting that AH7614 must enhance lipolysis by either inhibiting ligand independent FFA4 signalling, or FFA signalling that does not require the FFAs to be released from the cell. This study will help establish how FFA4 targeting therapeutics could be used to treat adipocyte dysfunction.

## Introduction

Adipose tissue is increasingly recognised for its important and diverse roles controlling metabolic health. While the function of adipose tissue and adipocytes in energy storage has long been understood, in recent years the broader importance of the tissue in glucose and lipid metabolism, as well as in cellular communication and signalling has been emerging.^1^ Dysfunction of adipose is well known to contribute to dyslipidaemia and glucose resistance, serving as an important driver of conditions like type 2 diabetes (T2D) and non-alcoholic fatty liver disease (NAFLD).^2^ Obesity is the main cause of adipocyte dysfunction and is characterised by increased adipocyte mass, caused either by hypertrophy of existing adipocytes or by adipogenesis leading to the development of new adipocytes.^3^ Considering the ever growing health burden linked to obesity and its associated metabolic disorders, there is a clear need to better understand how adipocyte dysfunction could be pharmacologically regulated.

Metabolite sensing G protein coupled receptors (GPCRs) are one important group of potential drug targets for the regulation of adipocyte function.^4^ This group of receptors includes a variety of GPCRs that have been shown to respond to intermediates of carbohydrate and lipid metabolism.^5^ In particular, a family of metabolite GPCRs that is activated by free fatty acids (FFAs) has received significant attention. The FFA family includes four members: FFA1 and FFA4 that are activated by medium and long chain fatty acids; and FFA2 and FFA3 that respond to short chain fatty acids.^6^ Of these, it is primarily FFA2 and FFA4 that have been found to be expressed in adipocytes.^7^ While there is good evidence that FFA2 controls aspects of adipocyte function including adipogenesis, lipolysis and glucose uptake,^8^ short chain fatty acids are produced primarily in the gut through fermentation of dietary fibre, making it difficult to study how pharmacological manipulation of FFA2 may interact with physiological regulation of the receptor using in vitro adipocyte cell models. In contrast, adipocytes serve an important role in the storage and release of medium and long chain fatty acids, suggesting in vitro cultured adipocytes could be a more physiologically relevant model to study FFA4 signalling. Supporting this are several recent studies that have suggested FFA4 function in adipocytes is regulated through local signalling pathways driven by lipolysis of stored fatty acids.^9, 10^ Therefore, cultured in vitro adipocytes provide an important model to explore how pharmacological manipulation of the FFA4 receptor may interact with physiological signalling mediated by FFAs produced within the adipocytes themselves.

The FFA4 receptor is a metabolite GPCR activated primarily by medium and long chain polyunsaturated fatty acids.^11, 12^ The receptor has historically been associated mainly with Gα_q/11_ signal transduction and recruitment of arrestin proteins, while more recent studies have increasingly also recognised its ability to signal through Gα_i/o_ pathways.^6^ As a result, early studies exploring FFA4 function in adipocytes focused on a role for the receptor enhancing insulin stimulated glucose uptake, as this is expected to be mediated by Gα_q/11_.^13^ However, relatively few follow up studies have reproduced this finding and the importance of FFA4 in either insulin dependent or insulin independent glucose uptake remains unclear. In contrast, the increased focus of FFA4-Gα_i/o_ signalling has led to interest in how this receptor may serve as a negative regulator of lipolysis,^9, 10, 14^ at least in part mediating a well-known negative feedback loop where fatty acids released during lipolysis inhibit the lipolytic pathway.^15^ In addition, FFA4 has also been shown to have a role in adipogenesis. Knockdown of FFA4 inhibits a murine 3T3-L1 model of adipogenesis,^16^ as does inhibition of the receptor in C3H10T1/2 murine pluripotent stem cells.^17^ However, despite evidence for the importance of FFA4 in adipocytes, to date a clear picture of how FFAs produced by adipocytes interact with pharmacological regulators of the receptor to influence adipogenesis, lipolysis and glucose uptake remains elusive.

Based on the view that FFA4 agonism may have benefit in the treatment of various metabolic disorders,^18^ a substantial number of small molecular agonists of this receptor have been developed.^19^ The best characterised of these is TUG-891, a potent FFA4 selective agonist that has been employed widely to study FFA4 pharmacology, structure and function.^6, 20–23^ In contrast, few FFA4 inhibitors or antagonists have been described and only a single compound, AH7614,^24^ is widely used. This compound has been shown to be a negative allosteric modulator of FFA4, exhibiting complex probe dependant pharmacology.^17^ However, some aspects of the pharmacology of AH7614, in particular whether this compound serves as an inverse agonist able to inhibit constitutive FFA4 signalling, remain unclear.

In the present study we have employed two FFA4 pharmacological tool compounds, TUG-891 and AH7614, to define the function of this receptor in mouse 3T3-L1 and human Simpson-Golabi-Behmel Syndrome (SGBS) adipocyte cell models. We first determine key signalling pathways and pharmacology associated with these ligands, showing that AH7614 acts as an allosteric inverse agonist of the receptor. We then establish how the pharmacological profile of the ligands may influence key adipocyte functions including adipogenesis, lipolysis and glucose uptake. Importantly, we find that while agonism of FFA4 has only modest effects on adipocyte function, AH7614 regulates each aspect of adipocyte function, either through inhibition of constitutive FFA4 activity or by inhibiting signalling in response to FFAs produced by the adipocytes themselves.

## Methods

### Materials

The FFA4 ligands, 4-[(4-Fluoro-4’-methyl[1,1’-biphenyl]-2-yl)methoxy]-benzenepropanoic acid (TUG-891) and 4-Methyl-*N*-9*H*-xanthen-9-yl-benzenesulfonamide (AH7614) were purchased from Tocris. The long chain fatty acid FFA4 agonist, α-linolenic acid (aLA) was purchased from Merck.

### Cell Culture

3T3-L1 cells were maintained in Dulbecco’s modified Eagle’s medium (DMEM) supplemented with 10% new-born calf serum (NCS) and penicillin/streptomycin. Cells were maintained at 37°C in a humidified cell culture incubator with 10% CO_2_ and subcultured before reaching confluency. For all experiments, 3T3-L1 cells were used at no more than passage eleven. Simpson-Golabi-Behmel Syndrome (SGBS) cells were as described previously.^25^ SGBS cells were maintained in DMEM/F12 supplemented with 1.7 mM D-Pantothenic acid, 3.3 mM biotin, penicillin/streptomycin, and 10% non-heat-inactivated foetal bovine serum (FBS). Cells were cultured at 37° C and 5% CO_2_ and used before passage five for all experiments.

Flp-In T-REx 293 cells engineered to express mouse FFA4 tagged with an enhanced yellow fluorescent protein were as described previously.^20^ Flp-In T-Rex 293 cells were maintained in DMEM supplemented with 10% FBS, 5 μg/ml blasticidin, penicillin/streptomycin and 200 μg/ml hygromycin B and maintained at 37° C and 5% CO_2_. HEK293T cells were maintained in DMEM supplemented with 10% FBS and penicillin/streptomycin and cultured at 37° C and 5% CO_2_.

### Adipogenic differentiation

To differentiate 3T3-L1 cells to adipocytes, cells were plated and grown to confluency and maintained for a further 48 h post-confluence in normal growth medium. To initiate differentiation, medium was replaced with DMEM containing 10% FBS, 0.5 mM 3-isobutyl-1-methylxanthine (IBMX), 5 μM troglitazone, 0.25 μM dexamethasone and 1 μg/ml insulin and cells cultured for 3 days. The medium was then replaced with DMEM supplemented with 10% FBS, 1 μg/ml insulin and 5 μM troglitazone before culturing the cells for a further 3 days. The medium was then replaced with DMEM containing 10% FBS and penicillin/streptomycin. Mature adipocytes were used between day 8 and 12 following initiation of differentiation.

SGBS cells were differentiated to adipocytes as previously described.^26^ Cells were seeded on collagen-coated plates and grown to near confluence. To initiate differentiation, cells were washed in Phosphate Buffered Saline (PBS) and medium changed to serum-free growth medium containing differentiation cocktail of 10 µg/ml apo-Transferrin, 100 nM human insulin, 2 nM triiodothyronine (T3), 100 nM dexamethasone, 500 µM IBMX and 1 µM rosiglitazone (day 0). On day 4, medium was changed to serum-free growth medium containing maintenance cocktail of 10 µg/ml apo-Transferrin, 10 nM insulin and 10 nM dexamethasone. Maintenance medium was replaced on day 8 and day 12, and adipocytes used on day 14.

### Oil Red O staining and quantification

Differentiated 3T3-L1 or SGBS cells were washed with PBS before fixing in 10% formalin for 30 minutes at room temperature. Cells were washed with PBS followed by 60% isopropanol before incubating for 15 min in 60% isopropanol containing 0.6% oil red O (ORO). The staining solution was discarded and cells washed again with 60% isopropanol followed by a wash with water. After drying, the stained cells were imaged using an EVOS FL AUTO 2 imaging system (ThermoFisher) fitted with a colour camera. Following imaging, quantification was carried out by first washed the cells twice with 60% isopropanol then extracting ORO stain in 100% isopropanol for 5 minutes. The isopropanol with extracted ORO was then transferred to a clear 96 well plate to measure absorbance at 492nm using a PHERAstar FS microplate reader (BMG LabTech).

Differentiated SGBS cells were ORO stained as described above. To control for cell number, before imaging, SGBS cell nuclei were stained by incubating with 10 µg/mL Hoescht 33342 (Invitrogen) for 15 minutes in the dark and washed. After imaging, Hoescht stain was quantified by measuring fluorescence intensity (ex: 360 nm; em: 460 nm) on a CLARIOstar plate reader (BMG LabTech). ORO was quantified as described and expressed as a ratio with fluorescence intensity for each well.

### qRT-PCR

RNA from 3T3-L1 or SGBS cells was prepared using an RNeasy kit (Qiagen), according to the manufacturer instructions. For SGBS cells, the on column DNAse I digestion protocol was used to eliminate genomic DNA, while for 3T3-L1 cells, isolated RNA was treated with DNAse I for 15 min. For cDNA synthesis, extracted RNA was reverse transcribed using a high-capacity cDNA reverse transcription kit with RNase inhibitor (Applied Biosystems) using random hexamer primers. Quantitative PCR reactions were then set up using a fast SYBR green master mix (ThermoFisher), with gene specific primers and cDNA. Real-Time PCR reactions were then carried out using QuantStudio 5 Real-Time PCR System (ThermoFisher) using the fluorescence channel for SYBR Green. To quantify expression, Ct values recorded from the reactions were analysed with the 2^-ΔΔCt^ method, using housekeeping genes hypoxanthine-guanine phosphoribosyl transferase (HPRT) for 3T3-L1 cells and transcription factor Sp1 for SGBS cells. Undifferentiated cells were used for each as the reference sample. Data were therefore expressed as a fold expression of that observed in undifferentiated cells.

### Western Blot

3T3-L1 adipocytes with indicate treatments were lysed in with ice cold RIPA Buffer (50 mM Tris, 150 mM NaCl, 2 mM MgCl_2_, 1% Triton, 0.5% sodium deoxycholate (w/v), 0.1% (w/v) SDS, 1 mM DTT, 50 units/ml Benzonase) for 30 min. The resulting lysate was then centrifuged at 21,910 x g for 3 min at 4° C and supernatants stored at −80° C until use. Cell lysates were mixed 1:1 with 2X Laemmli sample buffer (312.5 mM Tris-base, pH 6.8, 10% w/v SDS, 50% v/v glycerol, 250 mM DTT and 0.01% bromophenol blue) and heated to 60°C for 10 min. Samples were then resolved using sodium dodecyl sulphate polyacrylamide gel electrophoresis (SDS-PAGE), before transferring to a nitrocellulose membrane. Protein transfer was observed with Ponceau red, before membranes were washed with TBS (20 mM Tris-HCl, pH 7.5 and137 mM NaCl). Membranes were then incubated in TBS with 5% (w/v) milk powder for 60 min at room temperature to block non-specific binding sites. The indicated primary antibodies: rabbit anti-GLUT4,^27^ rabbit anti-phosphoAKT (Cell Signaling Technology) were then incubated, overnight at 4°C in blocking buffer. Following overnight incubation in primary antibody, nitrocellulose membranes were washed three times for 5 min with TBS containing 0.1% Tween 20 (TBST). Membranes were then incubated with an IRDye 800CW Donkey anti-Rabbit IgG (LICOR Biotechnologies) diluted in blocking buffer for 1 h at room temperature with shaking. Membranes were washed three times for 5 min with TBST followed by one wash with TBS for 5 min and finally one wash with H_2_O for 5 min. Blots were then imaged using a LICOR Odyssey SA scanner using the 800 nm laser.

### Glucose Uptake

3T3-L1 fibroblast were plated in 12-well plates and differentiated to adipocytes. The adipocytes were then washed once with serum-free DMEM then incubated with serum-free DMEM for 2 hours at 37° C. The cells were transferred to a hot plate maintained at 37° C and washed twice with Krebs-Ringer-phosphate buffer (128 mM NaCl, 1.25 mM CaCl_2_, 4.7 mM KCl, 5.0 mM NaH_2_PO_4_, 1.25 mM MgSO_4_). Cells were then treated with the indicated FFA4 ligand prior to the addition of insulin concentrations for 20 minutes. Thereafter, [^3^H]deoxyglucose/deoxyglucose solution was added to achieve a final concentration 60 μM deoxyglucose and 0.30 µCi/well and quickly mixed. After 3 minutes the reaction was terminated by quickly flipping the plates to remove isotope and dipping the plates in ice-cold PBS three times to stop the reaction. The plates were then allowed to dry for at least 30 minutes before the cells were solubilized overnight with 1% (v/v) Triton X-100. The solubilised material was then collected in scintillation vials with 5 ml scintillation fluid. A Beckman Multi-Purpose scintillation counter LS6500 was then used to quantify tritium in the sample.

### Lipolysis

3T3-L1 cells were grown and differentiated for 10 days. Cells were then washed twice with HBSS containing 25 mM glucose and 1% fatty acid-free bovine albumin serum, then cells were incubated with the same buffer either in the presence or absence of isoprenaline (1 nM) at 37°C for 1 h. Following the incubation, 50 μl of cell supernatant was transferred to a 96-well plate and mixed with 50 μl of free glycerol reagent (Merck). Plates were then incubated at 37° C for 15 min before absorbance at 540 nm was measured using a PHERAstar FS microplate reader (BMG LabTech). A standard glycerol curve was generated using a glycerol standard (Merck) and used to interpolate final glycerol concentrations.

### Ca^2+^ mobilisation

Flp-In T-REx cells engineered to express mFFA4-eYFP were seeded poly-D-lysine coated 96 black clear bottom plates and cultured until confluent. Cells were then induced to express the mFFA4 construct by adding 100 ng/ml doxycycline (dox) to the culture medium 24 h prior to the experiment. On the day of the assay, cells were incubated with the Ca^2+^ sensitive dye Fura-2 AM (1.5 μM) in culture medium at 37 °C for 45 minutes. The cells were then washed twice with Hanks’ Balanced Salt Solution containing 20 mM HEPES (HBSS) and incubated in HBSS at 37° C for 15 min. For experiments testing antagonists, the indicated concentration of antagonist was added before this 15 min incubation. The Ca^2+^ responses were then measured using a FlexStation^TM^ II microplate reader (Molecular Devices), monitoring Fura2 fluorescence (510 nm emission resulting from either 340 or 380 nm excitation) over a 90 second time period for each well. Fura2 ratios of 340/380 were calculated and the ratio obtained before the addition of test compound subtracted from the maximal ratio obtained over the 90 s measurement to calculate the overall Ca^2+^ response.

### TRUPATH G Protein Disassociation

G protein disassociation was assessed using bioluminescence resonance energy transfer (BRET) based biosensors. HEK293T cells were plated into 100 mm cell cultured dishes and grown to 60-70% confluency. Cells were then co-transfected with plasmids encoding mFFA4 tagged at its C terminal with an HA tag,^28^ and with a TRUPATH biosensor: consisting of plasmids encoding a Gα protein tagged with Renilla luciferase (Rluc), and the corresponding Gβ and a Gγ subunits as previously described for each Gα.^29^ The four plasmids were transfected in a 1:1:1:1 ratio using polyethyleneimine (PEI), with 30 μg of PEI and 5 μg of total plasmid DNA per dish. After transfection, cells were cultured for 24 h before subculturing into poly-D-lysine coated white 96 well plates at 50,000 cells/well. Cells were then cultured for a further 24 h before use. On the day of the assay cells were washed twice with HBSS then incubated in HBSS at 37° C for 30 min. The Rluc substrate Prolume Purple (NanoLight Technologies) was then added to a final concentration of 5 μΜ and the indicated concentrations of test compound added. Cells were incubated for 5 min before reading luminescent emission at 525 and 385 nm using a CLARIOStar microplate reader (BMG Labtech). BRET ratios were recorded as the ratio of 525 emission divided by 385 emission, then expressed as a fold of the BRET ratio obtained in vehicle treated cells.

### Arrestin-3 Recruitment

HEK 293T cells were plated into 100 mm dishes and cultured until they reached 60-70% confluency. Cells were then co-transfected with plasmids encoding: mFFA4-HA, arrestin-3 fused at its N terminal to Nanoluciferase (Nluc) and mNeonGreen (mNG) containing a CAAX lipid modification domain. A total of 5 μg of plasmid DNA was mixed with 30 μg of PEI per dish. Cells were then cultured for 24 h before subculturing at 50,000 cells/well in poly-D lysine coated white 96-well plates. Plates were cultured for a further 24 hours prior to the experiment. On the day of the experiment, cells were washed twice with HBSS and then incubated in HBSS for 30 min at 37° C. The Nluc substrate NanoGlo (Promega, N1110) was then added to a final 1:800 dilution and cells incubated in the dark for 10 min at 37° C. Bioluminescent emissions at 475 and 535 nm were then measured at regular intervals using a PHERAStar FS microplate reader (BMG Labtech) for 5 min before the indicated test compounds were added. Bioluminescent emission measurements were then continued for a further 30 min. Raw BRET ratios were taken as 535/475 nm emission ratio. Raw BRET ratios were then corrected for the BRET ratio obtained before the addition of test compound, followed by subtracting the response recorded from vehicle treated cells to obtain the NetBRET response. All data reported are the maximal change in BRET ratio observed over the 30 min experiment.

### Conditioned Medium and conditioned lipolysis experiments

For conditioned medium experiments, 3T3-L1 cells were plated and differentiated as described above using DMEM containing no phenol red, supplemented with 10% FBS and penicillin/streptomycin (basal medium). Conditioned medium (CM) was collected from confluent cells before differentiation was initiated (day 0 control) or 4 or 8 days post initiation of differentiation from wells where cells had been incubated in medium with or without differentiation cocktail. Medium containing differentiation cocktail was also collected from wells with no plated cells as a further control (naïve medium). CM was stored at l720° C until used in the assay. For conditioned lipolysis, fully differentiated 3T3-L1 cells were washed with HBSS before incubating in HBSS containing the indicated treatments (isoprenaline or TUG-891), with or without 1% BSA for 45 min. The resulting buffer was then collected and stored at l720° C until used in the assay.

To test whether conditioned medium or lipolysis experiment buffer contained fatty acids able to activate FFA4, arrestin-3 recruitment assays were carried out as described above by co-transfecting HEK293T cells with BRET biosensor components with or without hFFA4-HA. On the day of the assay, cells were washed twice with HBSS and then incubated in HBSS for 30 min at 37° C. HBSS was removed and replaced with the indicated conditioned medium or buffer and NanoGlo substrate was added to a final dilution of 1:800. Cells were incubated at 37° C in the dark for 10 mins before bioluminescent emissions at 475 and 535 nm were measured on the PHERAstar FS plate reader (BMG Labtech). Raw BRET ratios were taken as 535/475 nm emission ratio.

### Data Analysis

All data are presented as mean ± SEM from a minimum of 3 independent experiments. Data analysis and curve fitting were carried out using Graphpad Prism 10 (Graphpad Software Inc). Statistical analyses were carried out using t-tests, one way- or two way- ANOVA depending on the number of groups being compared and in all cases p<0.05 was view as statistically significant.

## Results

### FFA4 is upregulated during adipogenesis

3T3-L1 cells are a widely used murine derived cell model employed to study both adipogenesis and mature adipocytes.^30^ The cells are fibroblasts when maintained in normal culture, but can be differentiated into adipocytes following treatment with an adipogenic cocktail including insulin, IBMX, dexamethasone and troglitazone. Adipogenesis is characterised by an accumulation in neutral lipid, visualised as a striking increase in oil red O (ORO) staining of lipid droplets when comparing undifferentiated and differentiated cells (**Figure 1A**). Quantification of this ORO staining suggests a greater than 20-fold significant increase (p<0.001) in lipid after differentiation (**Figure 1B**). Adipogenesis in 3T3-L1 cells can also be monitored by assessing changes in transcript expression of key adipogenic markers, and significantly increased expression of transcript for adiponectin (p<0.05), leptin (p<0.05), PPARγ (p<0.05) and the insulin dependent GLUT4 glucose transporter (p<0.01) are observed after differentiation (**Figure 1C**). In contrast, expression of the insulin independent glucose transporter, GLUT1 (p<0.05), is decreased (**Figure 1Cv**). To determine how 3T3-L1 cells can be used to study FFA sensitive GPCRs in adipocytes, changes in transcript level expression for each FFA receptor was assessed before and after differentiation (**Figure 1D**). When examining the two short chain FFA receptors, FFA3 expression was not significantly affected by adipogenic differentiation, while FFA2 was upregulated by approximately 200-fold. For the long chain FFA receptors, FFA1 shows a modest decrease in expression following differentiation, while FFA4 exhibits a striking 37,000-fold increase (p<0.01) in transcript expression. Given that FFA1 and FFA4 are known to respond to many of the same ligands,^6^ this high upregulation of FFA4 combined with a downregulation of FFA1, suggests 3T3-L1 cells are a particularly useful model to determine the effects of FFA4 ligands on adipocytes.

**Figure 1.**
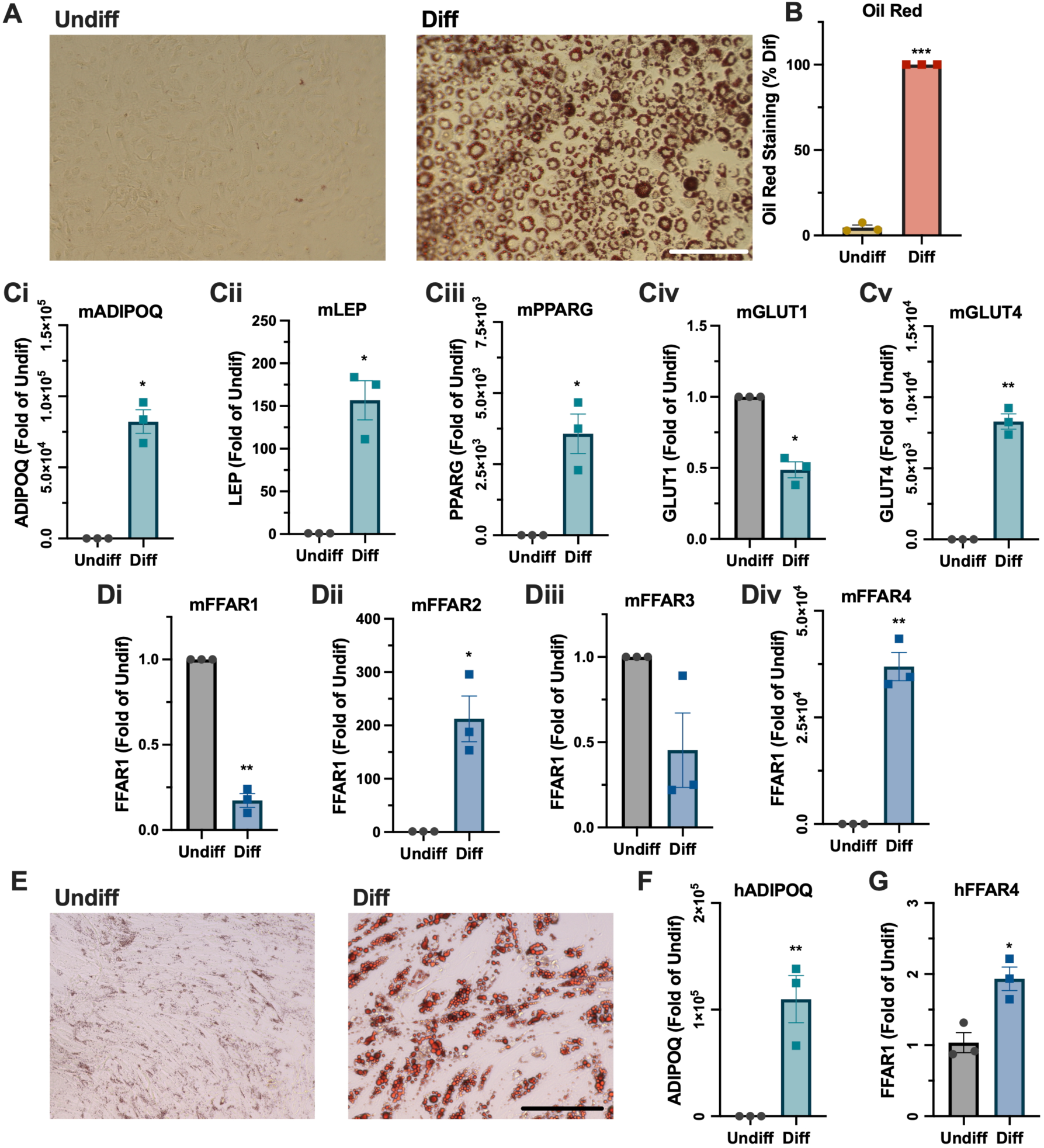
Mouse and human adipocyte cell models express FFA4 during differentiation. **A**. Brightfield images of 3T3-L1 cells stained with oil red O either before (Undiff) or after (Diff) a 9 day adipogenic differentiation. Scale bar is 200 μm. **B**. Quantification of oil red O staining from undifferentiated or differentiated 3T3-L1 Cells, n=3, *** p<0.001. Gene transcript expression of adipogenic markers (**Ci-Cv**) and free fatty acid receptors 1-4 (**Di-Div**) were assessed in 3T3-L1 cells using qRT-PCR. Data were analysed using the 2^-ΔΔCT^ method to show fold expression of undiff cells, using primers for HPRT as housekeeping control. N=3, ** p<0.01, * p<0.05. **E**. Brightfield images of SGBS cells stained with oil red O either without (Undiff) or after (Diff) adipogenic differentiation. Scale bar is 200 μm. Gene transcript expression of adipogenic marker ADIPOQ (**F**) and fatty acid receptor FFAR4 (**G**) were assessed by qRT-PCR. Data were analysed using the 2^-ΔΔCT^ method to show fold expression of undiff cells, using primers for Sp1 as housekeeping control. N=3 completed in duplicate, ** p<0.01, *** p<0.001.

To support findings in murine derived 3T3-L1 cells, SGBS cells are a human derived cell strain isolated from an infant believed to have Simpson-Golabi-Behmel Syndrome.^31^ These cells have many of the same properties as 3T3-L1 cells, showing a fibroblast like morphology when undifferentiated, but develop lipid droplets and accumulate neutral lipid following adipogenic differentiation (**Figure 1E**). SGBS cells also show a strong upregulation (p<0.01) of adiponectin transcript following differentiation (**Figure 1F**), comparable to the upregulation seen in 3T3-L1 cells (**Figure 1Ci**). Similarly, SGBS cells also upregulate (p<0.05) FFA4 transcript expression following differentiation (**Figure 1G**). Notably, the upregulation of FFA4 in SGBS cells is modest compared to what is observed in 3T3-L1 cells, however this may be due to relatively high FFA4 transcript expression in undifferentiated SGBS cells. Taken together, these findings suggest that SGBS cells represent a suitable human cell model to help understand how pharmacological regulation of FFA4 impacts adipocyte function.

### Complex FFA4 signal transduction

Before examining the function of FFA4 ligand in adipocyte models, we first aimed to characterise the signalling pathways of this receptor in heterologous expression systems. FFA4 has traditionally been associated primarily with Gα_q/11_ coupling but increasingly it is has also been linked to Gα_i/o_, while FFA4 interact with arrestins is also well established.^6^ To confirm activation of mouse FFA4 (mFFA4) leads to a Gα_q/11_ signalling, we found that both the synthetic FFA4 agonist, TUG-891, and a long chain FFA agonist, α-linolenic acid (aLA), produce concentration dependant Ca^2+^ responses in Flp-In T-REx 293 cells induced to express mFFA4 (**Figure 2A**). TUG-891 was found to be substantially more potent (pEC_50_= 6.5) than aLA (pEC_50_= 5.0). To specifically confirm Gα_q_ coupling of FFA4, a bioluminescent resonance energy transfer (BRET)-based TRUPATH G protein dissociation sensor,^29^ was employed in HEK293T cells expressing mFFA4 (**Figure 2B**). Again, these data show that both TUG-891 and aLA stimulate Gα_q_ activation, with TUG-891 again displaying higher potency: pEC_50_ of 6.3 for TUG-891 compared with 4.5 for aLA. To establish whether mFFA4 may also couple to Gα_i/o_, a Gα_i2_ TRUPATH sensor was used in HEK293T cells expressing mFFA4 (**Figure 2C**). Clear concentration dependant activation of Gα_i2_ was observed (TUG-891 pEC_50_ = 6.2; aLA pEC_50_ = 4.3), however the overall magnitude of BRET response did appear to be smaller for the Gα_i2_ sensor than it was when using the Gα_q_ sensor.

**Figure 2.**
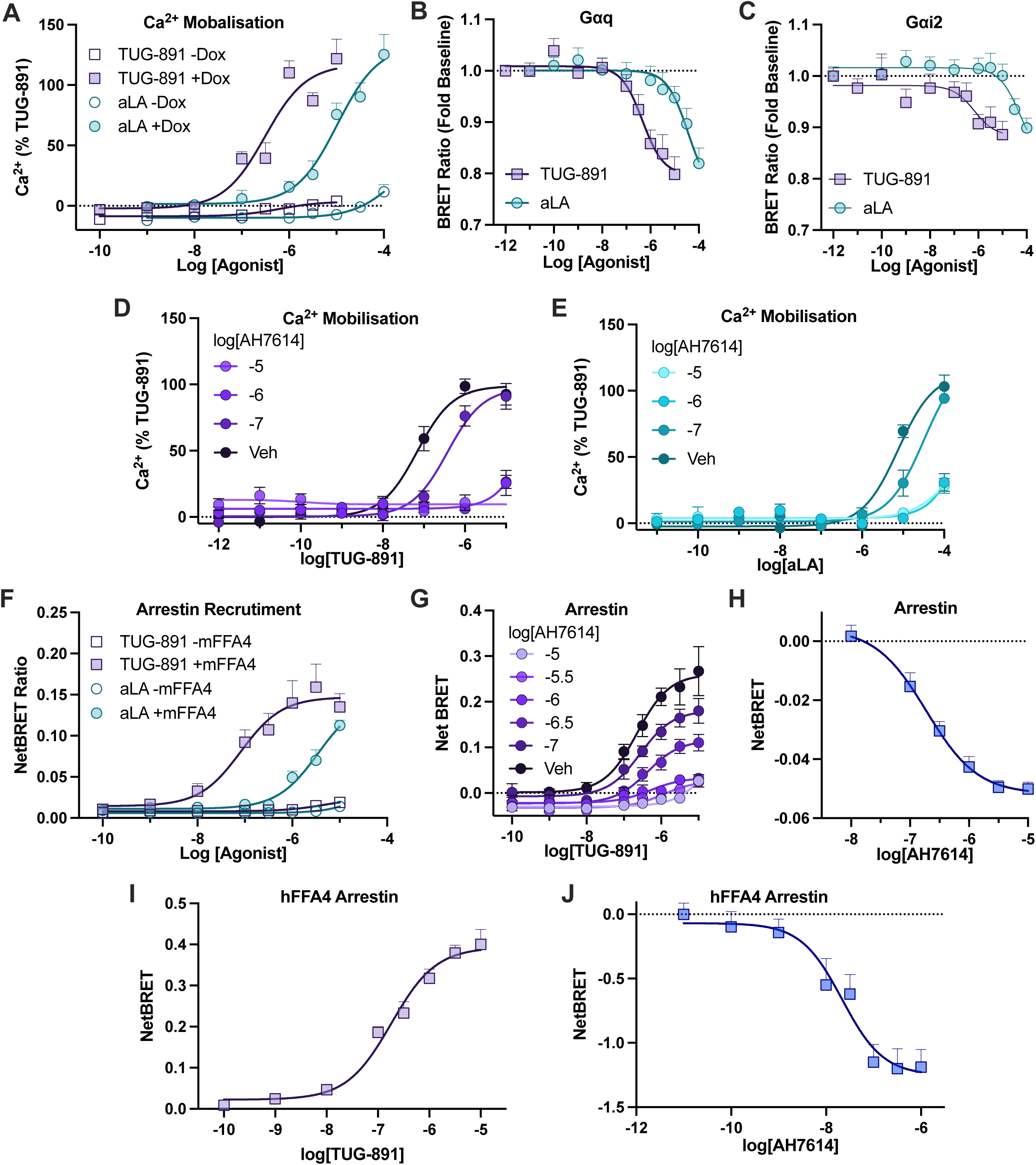
FFA4 allosteric antagonist, AH7614, behaves as an inverse agonist at both human and mouse FFA4. **A**. Ca^2+^ mobilisation responses to TUG-891 and aLA in Flp-In T-REx 293 cells that were either treated with dox (100 ng/ml), or not, to induce expression of mFFA4. N=3 completed in triplicate and expressed as a percent of the 10 μΜ TUG-891 response. G protein dissociation in response to TUG-891 and aLA was measured in HEK293T cells transfected with TRUPATH Gαq (**B**) and Gαi2 (**C**) biosensors. N=3 completed in triplicate. The ability of increasing concentrations of AH7614 to inhibit Ca^2+^ mobilisation responses to TUG-891 (**D**) and aLA (**E**) was assessed in Flp-In T-REx 293 cells induced to express mFFA4 with 100 mg/ml dox. N=3 in duplicate. Recruitment of arrestin-3 was measured in HEK cells expressing a bystander BRET arrestin-3 biosensor. **F**. Arrestin responses to TUG-891 and aLA treatment with or without expression of mFFA4. BRET responses are expressed as NetBRET above the signal obtained prior to the addition of ligand. N=3 in triplicate. **G**. Experiments measuring the ability of increasing concentrations of AH7614 to inhibit TUG-891 mediated arrestin recruitment to mFFA4. N=3 in duplicate. **H**. The effect of AH7614 alone on arrestin recruitment in cells expressing mFFA4 is shown. N=3 in triplicate. **I**. Data show arrestin-3 recruitment in HEK293T cells expressing a bystander BRET arrestin-3 sensor and hFFA4 and treated with increasing concentrations of TUG-891. N=3, **J**. Comparable experiments showing the arrestin response in cells expressing the arrestin-3 sensor and hFFA4 and treated with increasing concentrations of AH7614. N=3.

Having demonstrated that both agonists are able to activate FFA4-G protein signalling pathways, we next set out to confirm that FFA4-G protein signalling could be inhibited by the FFA4 allosteric antagonist AH7614.^17^ First we determined the effect of increasing concentrations of AH7614 on TUG-891 Ca^2+^ responses at mFFA4, observing that the inhibitor was able to effectively block almost all signalling (**Figure 2D**). Similarly, AH7614 effectively inhibited aLA mediated Ca^2+^ responses through mFFA4 (**Figure 2E**). These data suggest that AH7614 can be used as a tool to block G protein mediated signalling through mFFA4.

As FFA4 is also strongly associated with arrestin, a bystander BRET-based arrestin-3 biosensor was used,^32^ combining an Nluc tagged arrestin-3 with an mNeonGreen (mNG) fluorescent protein that was anchored to the cell membrane with a CAAX domain. This biosensor was transfected in HEK293T cells with or without mFFA4 DNA, and arrestin recruitment was assessed as an increase in BRET ratio (**Figure 2F**). Clear concentration dependent arrestin recruitment was observed for both TUG-891 (pEC_50_= 7.1) and aLA (pEC_50_= 5.5), only in cells that were expressing mFFA4. To determine if AH7614 could inhibit mFFA4 arrestin recruitment, concentration responses to TUG-891 were generated with increasing concentrations of AH7614 (**Figure 2G**). A notable suppression of the arrestin response was observed, manifest primarily as a reduction in maximal response. It was also notable that even at the highest concentrations of AH7614 tested, some recruitment appeared to remain, consistent with the view that this molecule is allosteric in its mode of action and often does not fully inhibit FFA4 signalling.^17^ Critically, it was also apparent that treating with increasing concentrations of AH7614 appeared to reduce the basal BRET signal even in the absence of TUG-891, and a more detailed analysis of the effect of AH7614 alone, clearly demonstrated a concentration dependent reduction in BRET (pIC_50_= 6.7) following AH7614 treatment alone (**Figure 2H**). These data suggest that the mFFA4 displays some ligand independent constitutive arrestin recruitment, and that AH7614 is able to act as an allosteric inverse agonist to suppress this activity.

Finally, we also aimed to confirm that AH7614 behaves in a similar way at human (h)FFA4. First, we confirmed that TUG-891 stimulated arrestin recruitment to hFFA4 could be measured using our Nluc-Arrestin-3/mNG-CAAX biosensor (**Figure 2I**). TUG-891 was able to potently stimulate arrestin recruitment with a pEC_50_ value of 6.7. Importantly, when testing the effect of AH7614 on arrestin recruitment to hFFA4 in the absence of agonist, this compound again behaved as an inverse agonist, resulting in a clear concentration dependent (pIC_50_ = 7.6) reduction in arrestin interaction (**Figure 2J**). In summary, our studies in heterologous cell systems confirm that TUG-891 is an effective FFA4 tool agonist at both mFFA4 and hFFA4, able to activate Gα_q/11_, Gα_i/o_ and arrestin pathways. In addition, we show that AH7614 is not only able to inhibit FFA4 agonist signalling, but also serves as an inverse agonist of ligand independent constitutive FFA4 signalling.

### Pharmacological inhibition of FFA4 suppresses adipogenesis

To test how pharmacological manipulation of the FFA4 receptor impacts adipogenesis, 3T3-L1 cells were differentiated in the presence of either vehicle, TUG-891, or AH7614 throughout the entire differentiation process, before accumulated lipid was stained with ORO (**Figure 3A**). Differentiation resulted in a clear increase in lipid in vehicle treated cells, with a comparable increase in TUG-891 treated cells, but reduced lipid staining was apparent in cells treated with AH7614. Quantification of this ORO staining confirmed the observation (**Figure 3B**), showing that while treatment with the FFA4 agonist, TUG-891, did not affect lipid accumulation (p>0.05), treatment with the FFA4 allosteric inverse agonist AH7614 significantly reduced (p<0.001) lipid staining to approximately 50% of the level observed in vehicle treated cells. A similar effect was seen when looking at lipid accumulation in human derived SGBS cells, where again a marked reduction in ORO staining was visible in cells differentiated in the presence of AH7614 (**Figure 3C**). Quantification of SGBS ORO staining indicated that lipid levels in these cells were reduce (p<0.01) to approximately 65% of control with AH7614 treatment. Together suggesting that while FFA4 agonism has little effect on adipogenesis, inverse agonism of the receptor inhibits adipogenesis in both human and mouse cell models.

**Figure 3.**
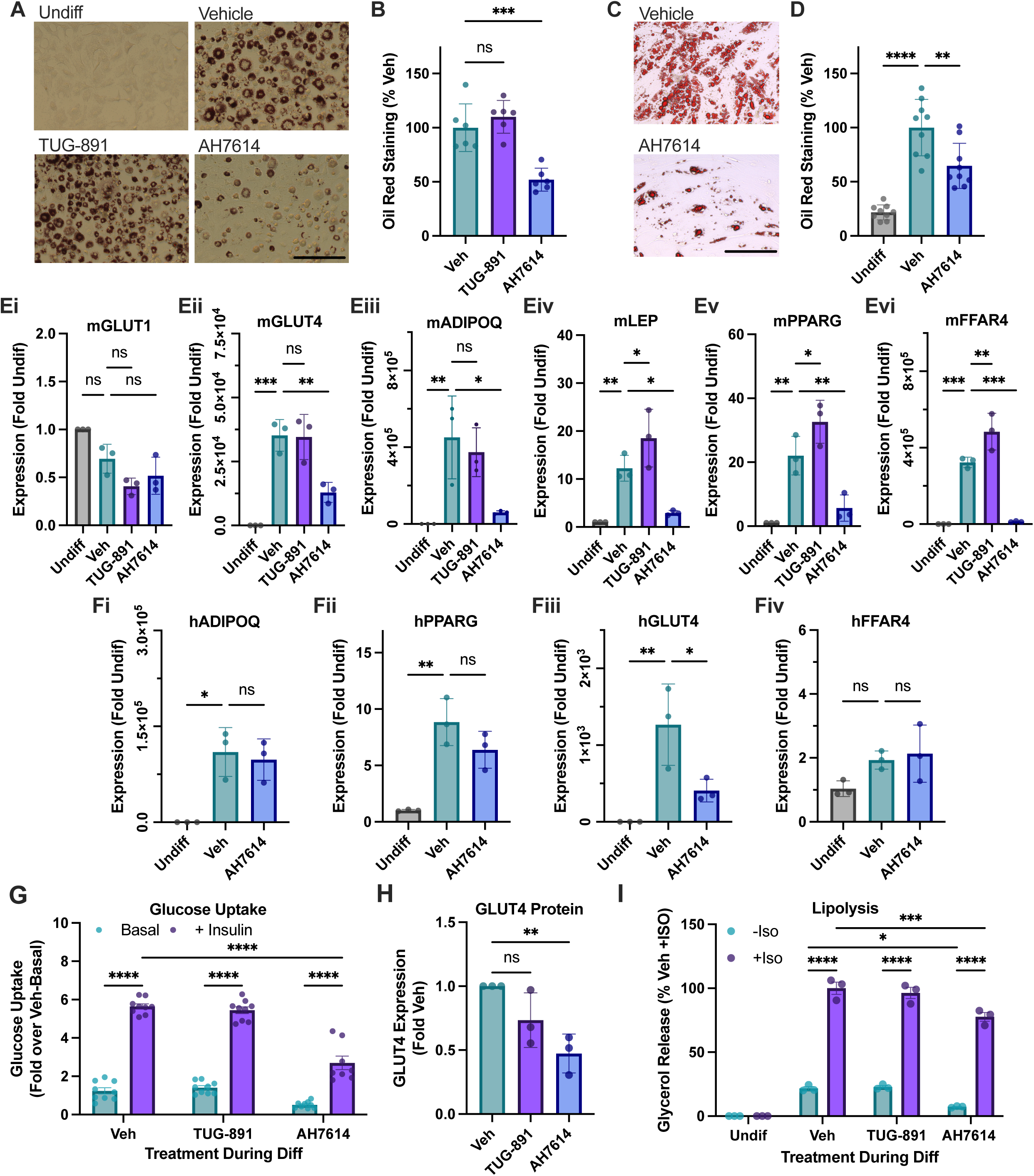
The FFA4 allosteric inverse agonist, AH7614, suppresses adipogenic differentiation in both mouse and human models. **A**. Brightfield images of oil red O stained 3T3-L1 cells that were either undifferentiated (Undiff), or exposed to adipogenic differentiation in the presence of 0.1% DMSO (Vehicle), TUG-891 (10 μM) or AH7614 (10 μΜ). Scale bar is 200 μm. Β. Oil red O staining was quantified and expressed as a percentage of the staining in vehicle treated cells. N=6, ***p<0.001. **C**. Brightfield images of oil red O stained SGBS cells exposed to adipogenic differentiation in the presence of either 0.1% DMSO (Vehicle) or AH7614 (10 μΜ). Scale bar is 200 μm. **D**. Quantification of oil red O staining from either undifferentiation SGBS cells, or from cells differentiated in the presence of vehicle or AH7614. N=3 in triplicate. Gene transcript expression of adipogenic markers (**Ei-Ev**) and FFAR4 (**Eiv**), assessed in 3T3-L1 cells by qRT-PCR following differentiation in the presence of vehicle, TUG-891, or AH7614. Data were analysed using the 2^-ΔΔCT^ method to show fold expression of undiff cells, using HPRT as housekeeping control. N=3, *p<0.05, **p<0.01, ***p<0.001. Gene transcript expression of adipogenic markers (**Fi-Fii**) and FFAR4 (**Fiv**), assessed by qRT-PCR in SGBS cells differentiated in the presence of vehicle or AH7614. Data were analysed using the 2^-ΔΔCT^ method to show fold expression of undiff cells, using Sp1 as housekeeping control. N=4 in triplicate, ** p<0.01, **** p<0.0001. **G**. 3T3-L1 cells were subjected to adipogenic differentiation in the presence of vehicle (0.1% DMSO), TUG-891 (10 μM), or AH7614 (10 μM). Differentiated adipocytes were then either treated or not (basal) with insulin for 20 min, before measuring uptake of [^3^H]deoxyglucose for 3 min. Glucose uptake is expressed as the fold of basal uptake in vehicle treated cells. N=3 In triplicate, **** p<0.0001. **H**. Protein level expression of GLUT4 assessed by western blot in 3T3-L1 cells differentiated in the presence of vehicle, TUG-891, or AH7614. GLUT4 protein level was first corrected for total protein on the blot and expressed as fold of the signal obtained in vehicle treated cells. N=3, **p<0.01. **I**. Glycerol release measured from either undifferentiated 3T3-L1 cells, or cells differentiated in the presence of vehicle, TUG-891, or AH7614. Cells were either untreated (-Iso) or treated with 1 nM isoprenaline (+Iso) for 1 h before measuring glycerol released. Data are expressed as a percentage of the glycerol level from vehicle differentiated cells treated with isoprenaline. N=3, *p<0.05, ***p<0.001, ****p<0.0001.

Next, to further confirm these effects of pharmacological manipulation of FFA4 in adipogenesis, 3T3-L1 cells were differentiated in the presence of either vehicle, TUG-891, or AH7614 throughout differentiation, before assessing transcript levels of key adipogenic markers by qRT-PCR (**Figure 3E**). Notably, AH7614 treatment significant suppressed expression of each adipogenic marker: GLUT4 (p<0.01), adiponectin (p<0.05), leptin (p<0.05), PPARγ (p<0.01), as well as supressing expression of FFA4 itself (p<0.001). Interestingly, although TUG-891 treatment did not result in enhanced lipid accumulation in the cells, it did increase expression of some adipogenic markers including: leptin (p<0.05), PPARγ (p<0.05), and FFA4 itself (p<0.01); suggesting that pharmacological FFA4 agonists may still have some role enhancing adipogenesis. Comparable experiments assessed the effect of AH7614 on adipogenic transcript expression in SGBS cells (**Figure 3F**), showing that despite effectively suppressing lipid accumulation in these cells, AH7614 had only modest effects on transcript expression of adipocyte markers. No effect was observed on adiponectin (**Figure 3Fi**) or FFA4 expression (**Figure 3Fiv**), while a modest but non-significant trend towards decreased PPARγ expression was seen (**Figure 3Fii**). Only GLUT4 expression was significantly suppressed (p<0.05, **Figure 3Fiii**), with AH7614 treatment reducing expression of GLUT4 transcript by 68%.

Having demonstrated that AH7614 suppresses adipogenesis, we next aimed to determine how differentiation of 3T3-L1 cells in the presence of this compound would affect mature adipocyte function. First, cells were exposed to either vehicle, TUG-891 or AH7614 throughout differentiation, before measuring the ability of the resulting mature adipocytes (without any FFA4 ligand present) to uptake [^3^H]2-deoxyglucose ([^3^H]2DG) in response to insulin treatment (**Figure 3G**). As expected, in cells differentiated in the presence of vehicle, insulin produces a clear increase in [^3^H]2DG uptake (p<0.0001). In cells differentiated with TUG-891, insulin also produced a significant increase in uptake (p<0.0001), which was essentially identical to the uptake observed in vehicle treated cells. In contrast, when cells were differentiated in the presence of AH7614, while insulin did still significantly increase [^3^H]2DG uptake (p<0.0001), the magnitude of uptake was significantly reduced (p<0.0001) to approximately 50% of the level observed in the vehicle differentiated cells. Considering we previously observed a reduction in GLUT4 transcript level with AH7614 treatment (**Figure 3Eii**), we hypothesised that the reduced glucose uptake was due to reduced expression of GLUT4 protein. To confirm this, we assessed GLUT4 protein expression by western blot, showing that there was indeed a significant (p<0.01) reduction in GLUT4 protein with AH7614 treatment during differentiation, to approximately 50% of the level observed in vehicle treated cells (**Figure 3H, Supplemental Figure 1**).

Finally, we assessed whether TUG-891 or AH7614 treatment during differentiation affected the ability of mature 3T3-L1 adipocytes to undergo lipolysis. Lipolysis was stimulated by treatment with a β-adrenoceptor agonist, isoprenaline (Iso), and measured through the release of glycerol into the assay buffer (**Figure 3I**). Notably, while no glycerol was release in undifferentiated cells either with or without Iso treatment, both the basal (-Iso), and stimulated (+Iso) levels of lipolysis increased following differentiation. Similar to what was observed in [^3^H]2DG uptake experiments, TUG-891 treatment during differentiation had no impact on either the basal or stimulated lipolysis when compared to the vehicle treated cells. In contrast, cells treated with AH7614 had reduced levels of both basal (p<0.05) and stimulated (p<0.001) lipolysis. Taken together, these data demonstrate that while pharmacological agonism of FFA4 has very little effect on adipogenesis, treatment with an FFA4 allosteric inverse agonist clearly suppressed adipogenic differentiation.

### AH7614 may inhibit autocrine regulation of FFA4 during differentiation

To better understand the mechanism by which TUG-891 and AH7614 influence adipogenic differentiation, 3T3-L1 cells were differentiated in the presence of DMSO vehicle, TUG-891 or AH7614 and lipid accumulation was quantified through ORO staining after 3, 6 and 9 days of differentiation (**Figure 4A**). After 3 days of differentiation no lipid accumulation was apparent in any of the conditions (**Figure 4Ai**), while clear accumulation was apparent by 6 days (**Figure 4Aii**). Interestingly, as we saw previously, while TUG-891 had no effect on lipid levels at day 9 (**Figure 4Aiii**), this FFA4 agonist did enhance lipid levels earlier in differentiation at day 6 (p<0.05). Perhaps suggesting that agonism of FFA4 may increase the rate of differentiation, even if it does not ultimately result in more lipid in the final mature adipocytes. AH7614 clearly supressed lipid levels at both day 6 (p<0.01) and at day 9 (p<0.01).

**Figure 4.**
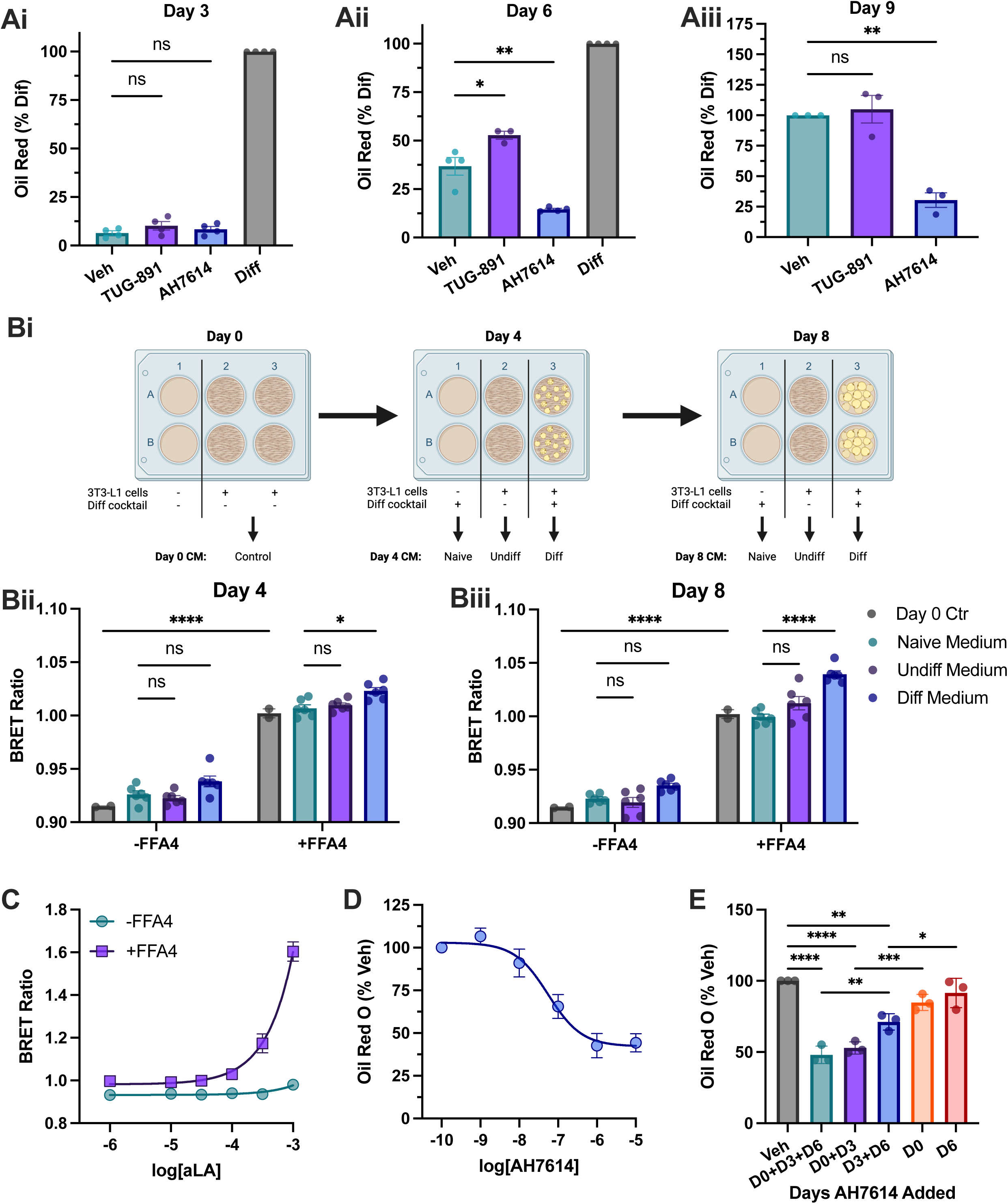
AH7614 inhibits FFA4 signalling to suppress differentiation throughout adipogenesis. **A**. 3T3-L1 cells were differentiated to adipocytes in the presence of 0.1% DMSO (Veh), TUG-891 (10 μM), or AH7614 (10 μM). Cells were fixed, stained with oil red O, and staining was quantified after day 3 (Ai), 6 (Aii), or 9 (Aiii) of differentiation. Staining is expressed as percent of oil red staining on day 9 vehicle control differentiated cells. N=3, *p<0.05, **p<0.01. **Bi.** Cartoon diagram outlining 3T3-L1 conditioned medium experiments. Conditioned medium from 3T3-L1 cells taken either 4 (**Bii**) or 8 (**Biii**) days into differentiation was tested for its ability to activate FFA4 activation in HEK293T cells expressing a bystander arrestin-3 biosensor. Experiments were conducted in cells expressing the biosensor and either transfected (+FFA4) or not (-FFA4) with FFA4 DNA. Day 0 medium was a control taken from cells that had not yet started differentiation, Naïve medium incubated in cell culture plates that had no cells in them, Undiff medium was from cells that had not been exposed to differentiation mediators, and Diff medium was from cells that were treated with differentiation mediators. N=3 in duplicate. **C**. A control experiment showing the concentration response for aLA to active a bystander arrestin-3 biosensor in cells expressing or not the FFA4 receptor. **D**. Quantification of oil red O staining in 3T3-L1 cells differentiated in the presence of increasing concentrations of AH7614. N=3, data are fit to a 3-parameter concentration response model. **E**. Quantification of oil red O staining in 3T3-L1 cells differentiated with AH7614 added only on the indicated days of differentiation. N=3, *p<0.05, **p<0.01, ***p<0.001, ****p<0.0001.

One possible explanation for why FFA4 antagonism inhibits differentiation, while agonism has little effect, could be that the FFA4 receptor is being activated by long chain FFAs that are being produced in the cells during adipogenesis.^33^ To explore this possibility, we conducted conditioned medium experiments where we took the culture medium (without phenol red) from differentiating 3T3-L1 cells and applied it to HEK293T cells expressing an FFA4 arrestin-3 biosensor (**Figure 4Bi**). These experiments aimed to determine if there were components of the culture medium able to activate the FFA4 receptor. For this, we compared conditioned medium taken from cells at day 0, with naïve medium that was incubated without cells, to medium incubated on cells with and without adipogenic differentiation mediators. Experiments were conducted with medium taken on day 4 (**Figure 4Bii**) or on day 8 (**Figure 4Biii**) of differentiation. When these conditioned media were applied to cells expressing the arrestin biosensor, but not the FFA4 receptor, no clear response to the biosensor was observed. However, when cells expressing FFA4 were used a clear increase in BRET was observed compared with the BRET recorded from cells not expressing FFA4. That was true even when comparing the control Day 0 medium or naïve medium, likely suggesting that this increase is due to constitutive ligand-independent FFA4 signalling. More importantly, when biosensor expressing cells were exposed to medium from both day 4 (p<0.05) and day 8 (p<0.0001), a significant increase in BRET was observed comparing the response to differentiation vs naïve medium, while no difference was observed between the undiff and naïve medium. Together these findings indicate that differentiation specifically results in the release of FFAs into the culture medium capable of activating FFA4. It was also notable that the BRET increase observed with the day 8 medium was greater than that produced using the day 4 medium, suggesting more FFAs were being released later in differentiation. To link the level of BRET response observed with the conditioned medium to concentrations of FFAs, a calibration experiment was conducted testing the ability of cells expressing the arrestin biosensor and FFA4 to respond to aLA diluted in naive culture medium (**Figure 4C**). It is notable that the potency of aLA is substantially lower in these experiments than in previous experiments that were not conducted in culture medium (**Figure 2F**); this is presumably due to the presence of FBS in the culture medium, and the fact that FFAs are well known to bind to serum proteins.^34^ Interpolation of the BRET values obtained in response to differentiation conditioned medium (**Figure 4B**), using the concentration-response curve for aLA (**Figure 4C**), suggests an FFA concentration of ∼100 μM in the diff medium on day 8, compared with ∼65 μM in the medium on day 4. These data suggest that AH7614 may be suppressing adipogenic differentiation by inhibiting autocrine FFA4 signalling in response to FFAs released from the cells as the adipocytes mature.

Next, in order to help confirm that AH7614 is mediating this effect specifically through inhibition of FFA4, a concentration response experiment was carried out. 3T3-L1 cells were differentiated for 9 days in increasing concentrations of AH7614 and differentiation was then assessed by ORO staining (**Figure 4D**). A clear concentration dependent decrease in lipid accumulation was observed, critically with a pIC_50_ (7.3) that matched well with the pIC_50_ values we observed for this compound inhibiting FFA4 signalling (**Figure 2H and 2J**). Finally, we also aimed to establish how the timing of AH7614 addition impacted its ability to suppress adipogenesis. 3T3-L1 cells were differentiated while adding AH7614 on day 0, on day 3 or on day 6 (or on combinations of these days) of differentiation then lipid accumulation was assessed by ORO staining on day 9 (**Figure 4E**). In these experiments it was notable that AH7614 tended to have a stronger effect when present early in differentiation. Notably, there was no difference in the effect of the compound (p>0.05) on lipid levels when only added on days 0 and 3, compared to when it was added on days 0, 3 and 6. In contrast, when AH7614 was added only on days 3 and 6, there was significantly less lipid accumulation (p<0.01). Adding AH7614 for only 3 days at the beginning or end of differentiation had no significant effect on lipid levels (p>0.05), suggesting 6 days of FFA4 inhibition is required to suppress adipogenesis. In summary, these finding support the view that FFA4 is responsible for the anti-adipogenic effects of AH7614 and that suppressing signalling earlier in differentiation has a greater impact.

### Agonism and inverse agonism of FFA4 regulates lipolysis in mature adipocytes

Having demonstrated that AH7614 suppresses adipogenic differentiation, we next aimed to determine what effect acute pharmacological regulation of FFA4 has on mature adipocytes. For this we first aimed to determine the effect of agonist, TUG-891, or allosteric inverse agonist, AH7614, on lipolysis. 3T3-L1 cells were differentiated before measuring both isoprenaline stimulated (+Iso) and basal (-Iso) lipolysis through the release of glycerol. Addition of TUG-891 was found to significantly reduce isoprenaline stimulated lipolysis at 10 μM (p<0.0001) and 30 μM (p<0.0001) concentrations (**Figure 5A**). In contrast, treatment with AH7614 significantly enhanced isoprenaline stimulated lipolysis at 10 μM (p<0.05) and 30 μM (p<0.0001) concentrations (**Figure 5B**). These findings are broadly consistent with previous work suggesting FFA4 may serve as an autocrine regulator of lipolysis.^9^ However, more recent studies have raised questions as to whether FFAs produced and released into the culture medium during lipolysis are sufficient to activate the FFA4 receptor at the cell surface.^10^ To explore this in more detail, we again carried out conditioned medium experiments where 3T3-L1 adipocytes were either treated with isoprenaline or vehicle in the same buffer used for lipolysis experiments, before transferring this buffer to HEK293T cells co-expressing FFA4 with an arrestin-3 bystander BRET biosensor (**Figure 5C**). Notably, in these experiments the buffer taken from cells treated with isoprenaline did not increase arrestin-3 recruitment to FFA4, suggesting the buffer did not contain sufficient FFA levels to activate the receptor (**Figure 5D**). Notably, control buffer that contained the FFA4 agonist, TUG-891, was able to produce a significant increase in arrestin recruitment. One possible explanation for the inability of isoprenaline treated lipolysis buffer to stimulate FFA4 activation could be the presence of BSA in the buffer. In lipolysis experiments BSA is required to accept the lipophilic fatty acids being released,^15^ but in doing so it also likely reduces the ability of the fatty acids to bind to and activate FFA4.^6^ Therefore, we repeated these experiments using lipolysis buffer where the BSA had been removed (**Figure 5D**). Again, conditioned medium from isoprenaline treated cells did not result in any increase in FFA4 activation, in this case likely because without BSA only very low levels of fatty acid will be released.^10^ However, it was notable that when BSA was removed the ability of the control buffer containing TUG-891 to activate arrestin recruitment was substantially improved (BRET ratio of 0.97 with BSA compared with 1.34 without). Overall, these finding confirm the ability of both an FFA4 agonist, TUG-891, and an allosteric inverse agonist, AH7614, to regulate lipolysis in mature adipocytes. However, our findings suggest that regulation of lipolysis by AH7614 is not dependent on inhibition of response to FFAs that have been released in an autocrine fashion into the culture medium.

**Figure 5.**
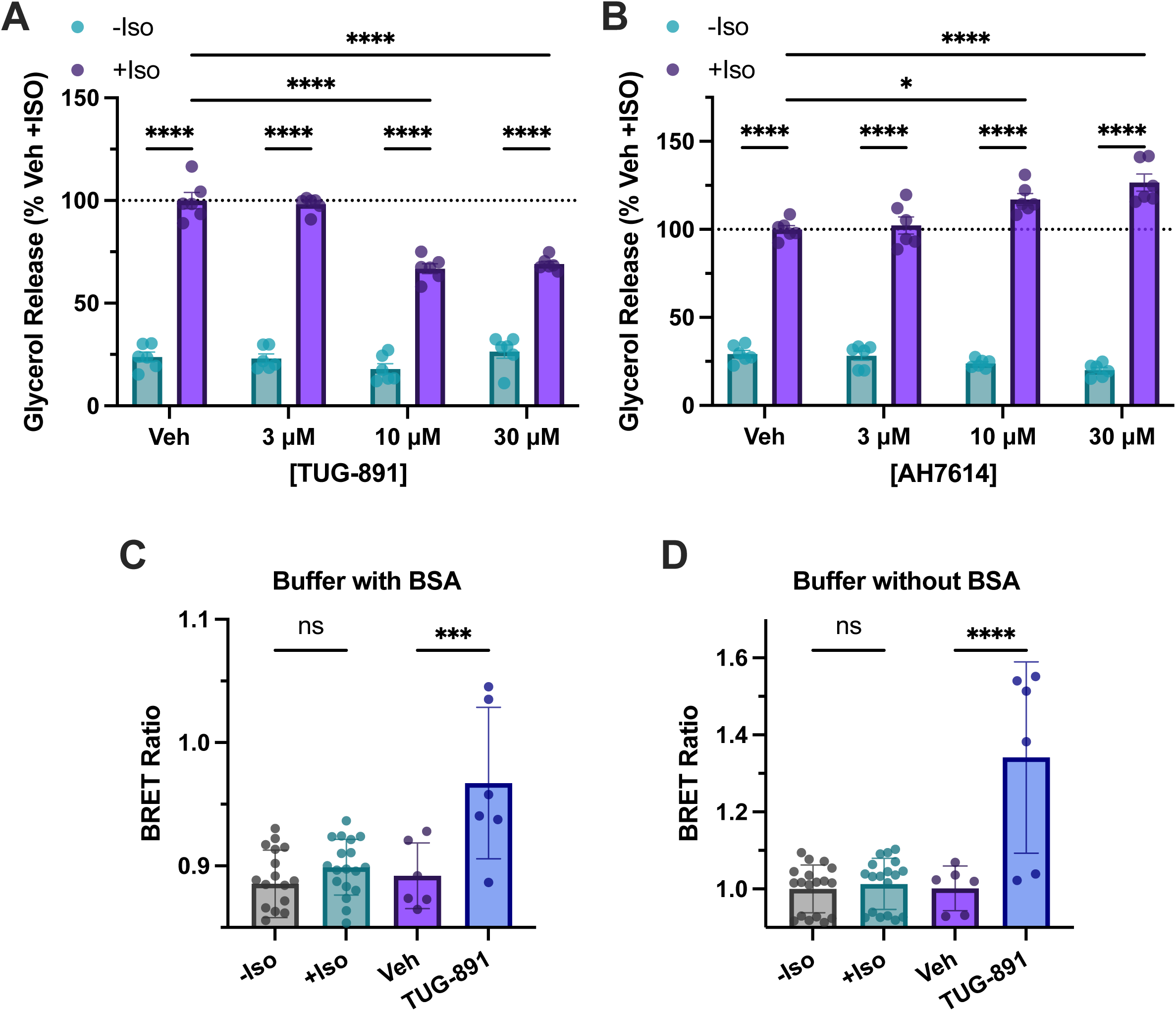
FFA4 regulates lipolysis in 3T3-L1 adipocytes. **A**. Differentiated 3T3-L1 adipocytes were treated with either 0.1% DMSO (Veh), or increasing concentrations of TUG-891 for 45 min. Cells where then either treated (+Iso) or not (-Iso) with isoprenaline (1 nM) for 1 h and glycerol released was quantified. Data are expressed as percentage of glycerol released in the +Iso/Veh condition. N=3 in duplicate, ****p<0.0001. **B**. Comparable experiments to those shown in **A**, but with increasing concentrations of AH7614. N=3 in duplicate, *p<0.05, ****p<0.0001. **C**. Conditioned medium experiments using lipolysis assay buffer applied to 3T3-L1 cells for 1 h either with (+Iso) or without (-Iso) isoprenaline; with 0.1% DMSO (Veh) or with TUG-891 (10 μM). This conditioned buffer was then applied to HEK293T cells co-expressing FFA4 and a bystander BRET arrestin-3 biosensor, and the BRET ratio was recorded after 5 min. N=3, ***p<0.001. **D**. Comparable experiments to those shown in **C**, but using a modified lipolysis assay buffer where BSA was removed. N=3, ****p<0.0001.

### FFA4 inverse agonism suppresses glucose uptake in adipocytes

Finally, we aimed to determine how acute pharmacological regulation of FFA4 influences glucose uptake in mature 3T3-L1 adipocytes. First, we assessed the effect of TUG-891 on basal, insulin independent, [^3^H]2DG uptake (**Figure 6A**). In these experiments, despite a robust glucose uptake in response to insulin, no significant effect on uptake was observed for any of the concentrations of TUG-891 tested (p>0.05). In contrast, when basal [^3^H]2DG uptake was measured following treatment with AH7614 (**Figure 6B**), significant reductions in basal uptake were observed at both 3 μΜ (37%, p<0.01) and 30 μM (36%, p<0.01) AH7614 concentrations. There were also notable trends towards decreased uptake at all other concentrations tested, including 0.3 μM (19%, p=0.15), 1 μΜ (19%, p=0.16) and 10 μM (22%, p=0.10). These data suggest that acute AH7614 treatment suppresses insulin independent [^3^H]2DG uptake in adipocytes.

**Figure 6.**
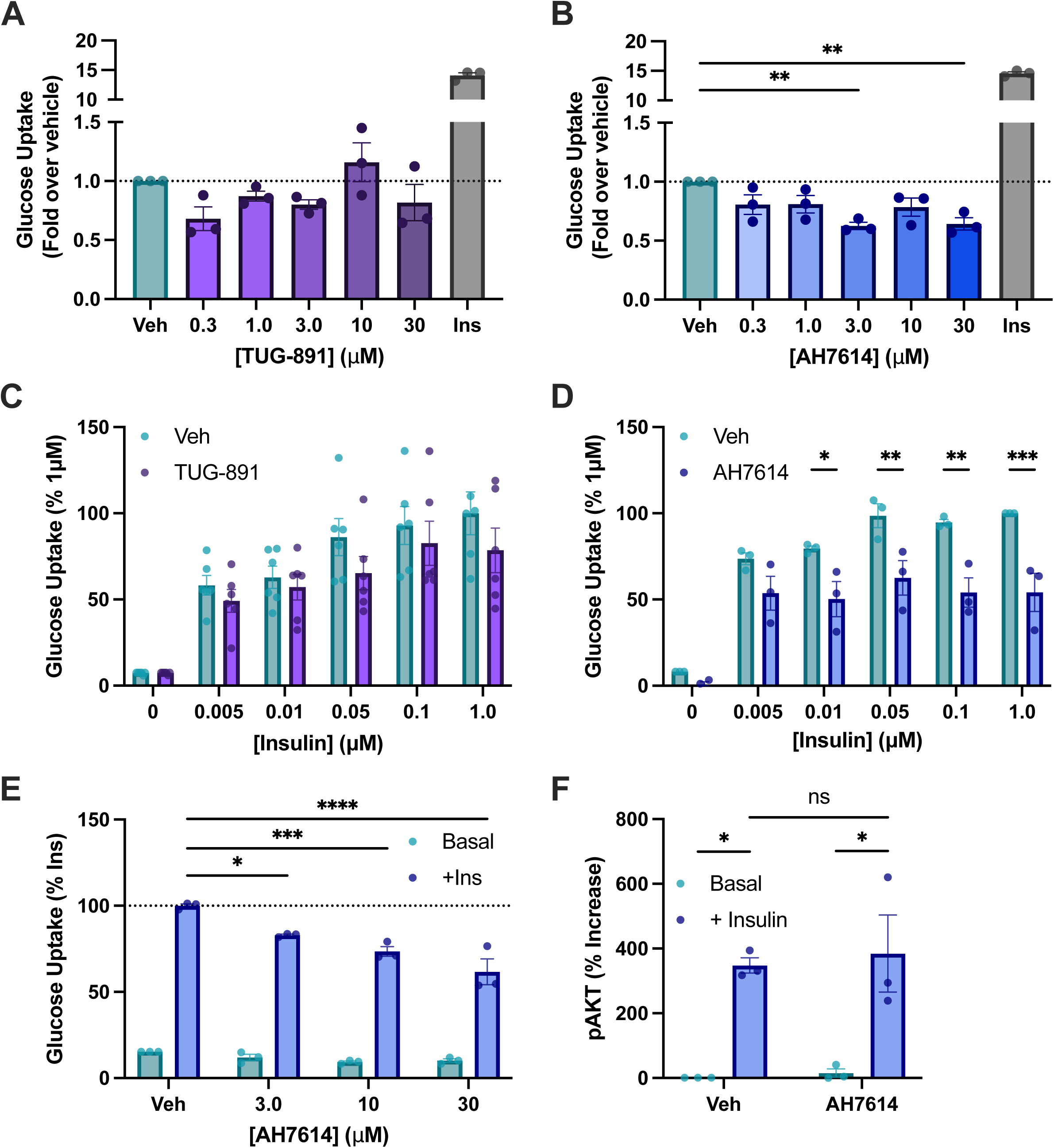
The FFA4 allosteric inverse agonist, AH7614, inhibits both insulin dependent and insulin independent glucose uptake. **A**. Uptake of [^3^H]deoxyglucose in differentiated 3T3-L1 adipocytes was measured for 3 min following a 45 min treatment with 0.1% DMSO (Veh) or with increasing concentrations of TUG-891. A control treatment with 1 μM insulin (Ins) for 20 min is included for reference. N= 3. **B**. Comparable experiments with increasing concentrations of AH7614. N=3, **p<0.01. **C**. Uptake of [^3^H]deoxyglucose in differentiated 3T3-L1 adipocytes in cells pre-treated for 30 min with either 0.1% DMSO (Veh), or TUG-891 (10 μM). Cells were then treated with increasing concentrations of insulin for 20 min, before measuring uptake over a 3 min period. Uptake was expressed as a percentage of that measured in Veh/1μM insulin treated cells. N=6 in triplicate. **D**. Comparable experiments but with cells treated with AH7614 (10 μM) for 20 min instead of with TUG-891, prior to the addition of insulin. N=3 in triplicate, *p<0.05, **p<0.01, ***p<0.001. **E**. Differentiated 3T3-L1 cells were pre-treated with either 0.1% DMSO (Veh), or with increasing concentrations of AH7614 for 20 min before treating 20 min, with or without (basal), 1 μM insulin. [^3^H]deoxyglucose was then measured over a 3 min period and expressed as a percentage of the uptake measured in veh/insulin treated cells. N=3, *p<0.05, ***p<0.001, ****p<0.0001. **F**. Phosphorylation of AKT was assessed by western blot from lysates of differentiated 3T3-L1 adipocytes pre-treated with either 0.1% DMSO (Veh) or AH7614 (10 μΜ), followed by 20 min treatment with or without 1 μM insulin. N=3, *p<0.05.

We moved on to determine if either TUG-891 or AH7614 affected insulin dependent glucose uptake in mature adipocytes. When measuring the uptake of [^3^H]2DG in response to increasing concentrations of insulin, we found that treatment with either DMSO vehicle or TUG-891 had not significant effect on the level of glucose uptake observed (**Figure 6C**). This was true, despite the fact that a clear concentration dependent uptake response was observed to increasing concentrations of insulin. In contrast, when comparable experiments were carried out using the FFA4 allosteric inverse agonist, AH7614, clear inhibition of insulin dependent uptake was observed at 0.01 (p<0.05), 0.05 (p<0.01), 0.1 (p<0.01), and 1.0 μM (p<0.001) insulin concentrations (**Figure 6D**). To explore this relation between AH7614 and insulin signalling further, the ability of increasing concentrations of AH7614 to inhibit [^3^H]2DG to 1 μM insulin was examined (**Figure 6E**). Again, not only did AH7614 clearly suppress insulin stimulated uptake, it did so in a concentration-dependant manner. Finally, to explore where in the signalling pathway AH7614 may be acting, phosphorylation of AKT was assessed in cells treated with insulin and or with AH7614 (**Figure 6F, Supplemental Figure 2**). As expected, insulin treatment robustly increased AKT phosphorylation, however this was unaffected by AH7614, suggesting that the effect of the compound must occur downstream of AKT phosphorylation. In summary, our glucose uptake experiments indicate that although FFA4 does have a clear role regulating uptake, pharmacological manipulation of this can be achieved using an FFA4 inverse agonist, but not with an FFA4 agonist.

## Discussion

Adipocyte dysfunction is a hallmark of many metabolic disorders, including obesity, NAFLD and T2D. This dysfunction can relate both to adipogenic development of new adipocytes and to the normal function of mature adipocytes.^1^ There is a clear need for new pharmacological interventions that can help restore normal adipocyte function. In the current study we have explored how pharmacological manipulation of the FFA4 receptor impacts adipogenesis and two key mature adipocyte functions: isoprenaline stimulated lipolysis and insulin stimulated glucose uptake. We show that while pharmacological activation of FFA4 with a synthetic agonist, TUG-891, tends to have only modest effects, inhibition of FFA4 using a negative allosteric modulator, that we show to also poses inverse agonism properties, AH7614, modulates all adipocyte functions tested. Importantly, because FFA4 is activated by long chain FFAs, which are produced and released in high levels from adipocytes, we demonstrate that some of AH7614 effects are likely due to inhibition of FFA/FFA4 signalling loops occurring in the adipocytes.

The FFA4 receptor is a GPCR that is traditionally associated with Gα_q/11_ signalling and arrestin recruitment, but that has received increased focus on Gα_i/ο_ signalling in recent years.^6^ Our findings on in heterologous systems generally support this view. We observed that FFA4 agonists stimulate Gα_q/11_ signalling through dissociation of Gα_q_ and intracellular Ca^2+^ mobilisation, while also robustly recruiting arrestin-3 to the cell membrane. Importantly, we also demonstrate that mFFA4 agonism stimulates dissociation of Gα_i2_, indicating that this receptor does also engage with Gα_i/o_ pathways. These observations informed our choice of adipocyte functional assays, as lipolysis is well known to be negatively regulated by Gα_i/o_ signalling,^35^ glucose uptake is enhanced by Gα_q/11_,^36^ and adipogenesis can be promoted through both Gα_i/o_ and Gα_q/11_ pathways.^37^

Our pharmacological characterisation of the only reported FFA4 antagonist, AH7614, demonstrated that not only does this compound inhibit agonist stimulated G protein and arrestin pathways, but it also acts as an inverse agonist, inhibiting ligand-independent, constitutive activity of FFA4. Importantly, we show that this property is observed at both human and mouse orthologs of FFA4, something particularly important given that there are substantial species ortholog variations in pharmacology for many of the FFA receptors.^38^ Since AH7614 has been shown to be a negative allosteric modulator,^17^ our findings demonstrate for the first time that AH7614 is actually an allosteric inverse agonist of FFA4. Constitutive activity of GPCRs occurs when the receptor adopts an active conformation in the absence of an agonist, and although it has been observed for some of the FFA receptors,^6^ it has not been widely reported for FFA4. It is however important to consider that FFAs (if mainly esterified) are present in all cells, and therefore it can be difficult to be sure if constitutive signalling from an FFA receptor is due to true ligand-independent constitutive activity, or to the receptor responding to endogenous fatty acids.^39, 40^ Future studies should aim to more directly explore the importance and significance of FFA4 constitutive activity.

Our observation that pharmacological inhibition of FFA4 during adipogenic differentiation suppresses adipogenesis in both human and mouse cell models is broadly consistent with previous work linking FFA4 to adipogenesis.^41^ Previous studies have used genetic knockdown in 3T3-L1 and other adipogenic cell models, finding that reducing FFA4 expression inhibits lipid accumulation and expression of adipogenic markers.^16, 42, 43^ Our experiments extend this work, not only by showing the effect can be achieved pharmacologically, but critically, also demonstrating that inhibition during adipogenesis translates to reduce mature adipocyte response to insulin and isoprenaline in glucose uptake and lipolysis experiments, respectively. This highlights the complexity and sometimes counterintuitive view that promoting adipogenesis could be beneficial in dysregulated adipose, as it may help restore normal metabolic function, including insulin responsiveness.^3^ More importantly, through conditioned medium experiments, our study has demonstrated that during adipogenesis 3T3-L1 cells are releasing FFAs into the culture medium that are sufficient to activate the FFA4 receptor. While previous studies have shown that this may be the case in mature adipocytes stimulated to induce lipolysis,^9^ it was unclear whether basal FFAs released during adipogenesis would be sufficient to activate the receptor. Our observation indicates that the role of FFA4 in adipogenic differentiation is to mediate a positive feedback autocrine signalling loop, where FFAs released feedback on the differentiating cells to further enhance differentiation.

One of the more unexpected outcomes of this study was the observation that TUG-891 treatment had only modest effects on both adipogenic differentiation and on glucose uptake. Previous work has suggested that FFA4 agonism does enhance glucose uptake,^13, 20^ and while studies on the effects of the FFAs themselves on adipogenesis have been mixed, potent synthetic FFA4 agonists like TUG-891 have tended to show pro-adipogenic effects.^42, 43^ It is however worth noting that it in previous studies on 3T3-L1 cells where TUG-891 did show an effect, a less robust adipogenic differentiation protocol was used lacking a thiazolidinedione PPARγ agonist.^42^ Given that thiazolidinediones are well known to enhance adipogenesis,^44^ part of the reason we do not see a strong effect to TUG-891 may be due to the fact we have included a thiazolidinedione during differentiation, perhaps making it difficult to further enhance the adipogenic process. Further complicating this situation is the fact that natural FFA agonists of FFA4 are activators of PPARγ themselves,^45^ and synthetic FFA4 agonists have also been shown to indirectly activate this transcription factor.^46^ The fact that in our hands AH7614 still inhibits adipogenic differentiation, even when using a robust protocol that includes the thiazolidinedione, reinforces the importance of FFA4 in adipogenesis.

Although agonism of FFA4 with TUG-891 did not affect glucose uptake in the mature adipocytes, inhibition of the receptor with the allosteric inverse agonist, AH7614 suppressed both basal uptake and insulin dependent uptake. This observation suggests that either ligand-independent constitutive activation of FFA4, or basal activation of FFA4 by endogenously produced FFAs must have some role in mediating the glucose uptake response. Treatment of mature adipocytes with AH7614 was also found to enhance isoprenaline stimulated lipolysis, again suggesting basal or constitutive FFA4 activation may play a role. Given one of the main products of lipolysis is FFAs, and previous work suggested that FFA4 functions in a negative autocrine feedback look to control lipolysis,^9^ we hypothesised that in this case AH7614 was likely inhibiting signalling due to FFAs that were released during lipolysis. Surprisingly, our conditioned medium experiments did not show that there were sufficient levels of FFA present in the buffer from our lipolysis experiments to activate the FFA4 receptor. This was true whether BSA was included in the buffer to accept the FFAs or not. These findings strongly suggest that AH7614 is not enhancing lipolysis by inhibiting FFA4 signalling mediated from FFAs that have been released into the culture medium. A recent study examining the role of FFA4 in lipolysis has found that in adipocytes the FFA4 receptor is not only expressed at the cell surface, but is also found on intracellular membranes lining the lipid droplets.^10^ More interestingly, the study demonstrated that instead of functioning in an autocrine signalling pathway to inhibit lipolysis, intracellular FFA4 receptors appear to be activated by fatty acids in an ‘intracrine’ manner, as the FFAs are being produced from the lipid droplets. This therefore suggests two possible mechanisms by which AH7614 enhances lipolysis: inhibiting constitutive ligand independent FFA4 activity or inhibiting intracrine FFA4 signalling. Given that AH7614 is a relatively lipophilic molecule and likely could cross the cell membrane, either of these two possibilities, or perhaps a combination of the two are plausible.

In summary, we have shown that pharmacological manipulation of the FFA4 receptor has profound effects on both adipogenic differentiation and mature adipocyte function. While we do observe some effect of FFA4 agonism, consistently we see that inhibition of the receptor has a greater impact. We show that FFA4 displays a background level of constitutive activity and propose that this fact, combined with naturally produce FFAs in adipocytes results in a high background activation of FFA4, which can therefore not be further activated by agonist treatment. In contrast, the inverse agonism properties of AH7614 mean that this compound is able to inhibit both constitutive and FFA mediated FFA4 signalling, explaining why AH7614 is able to modulate all FFA4 pathways in our adipocyte models. Together, our findings help to provide a deeper understanding of how this important receptor regulates adipocyte function and how it may be exploited therapeutically to address adipocyte dysfunction.

## Acknowledgments

This work was supported by an academy of medical science springboard award (BDH, SBF004\1033) and by a research scholarship from the University of Hafr Al Batin (WSA). The work was also supported by the EPSRC and SFI Centre for Doctoral Training in Engineered Tissues for Discovery, Industry and Medicine, Grant Number EP/S02347X/1 (EMD), and a Medical Research Scotland Studentship (LV and BDH, PHD-50838-2024).

## CRediT Author Contributions

Conceptualization (WSA, GG, BDH), Formal analysis (WSA, EMD), Funding acquisition (WSA, GG, BDH), Investigation (WSA, EMD, LV, MK, BD), Methodology (MW), Supervision (GG, BDH), Writing-original draft (BDH), writing review and editing (WSA, EMD, LV, MK, BD, MW, GG, BDH).

***Sup Fig 1.** Images of GLUT4 Western Blot*. **A**. Lysates collected from undifferentiated 3T3-L1 cells or from cells differentiated in the presence of 0.1% DMSO (Veh), TUG-8891 (10 μM) or AH7614 (10 μM) were used in a western blot with an anti-GLUT4 antibody. GLUT4 bands run as a smear due to glycosylation. **B**. Total protein stain of the blot shown in **A**. Quantification was determined as the total intensity of staining in the GLUT4 blot divided by total intensity in the total protein stain.

***Sup Figure 2.** Images of pAKT Western Blot*. Lysates were collected from differentiated 3T3-L1 cells that were first pre-treated with 0.1% DMSO vehicle or AH7614, followed by a 20 min treatment with or without 1 μM insulin. For the western blot 20 μg of protein was used and blots either probed with an anti-phosphoAKT (top) antibody or stained with a total protein stain (bottom). Quantification was determined as the intensity of the pAKT band divided by total intensity in the total protein stain.

