## Supplementary figures and images for "Inverse Agonism of the FFA4 free fatty acid receptor controls both adipogenesis and mature adipocyte function"

### Supplemental Figure 1

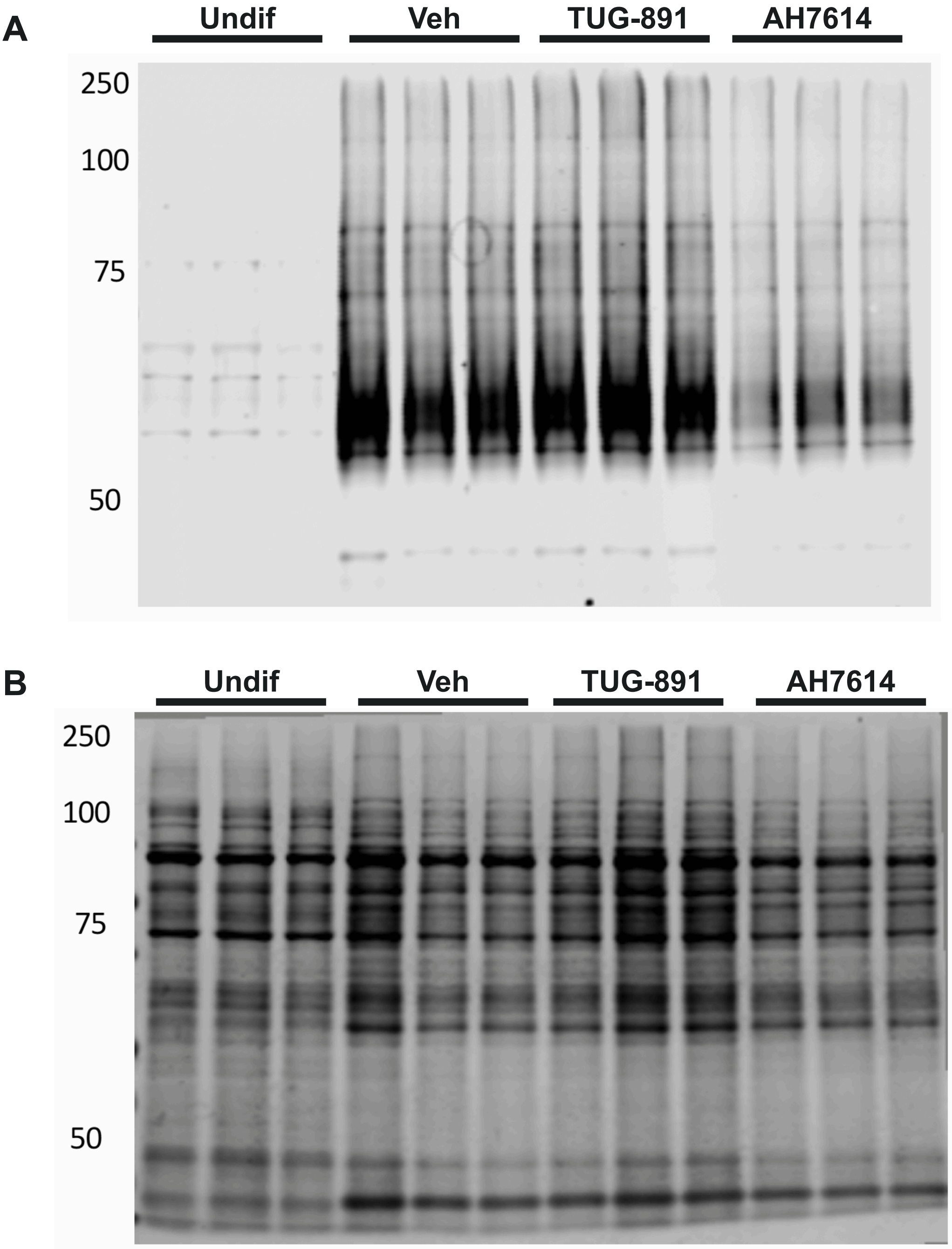

### Supplemental Figure 2

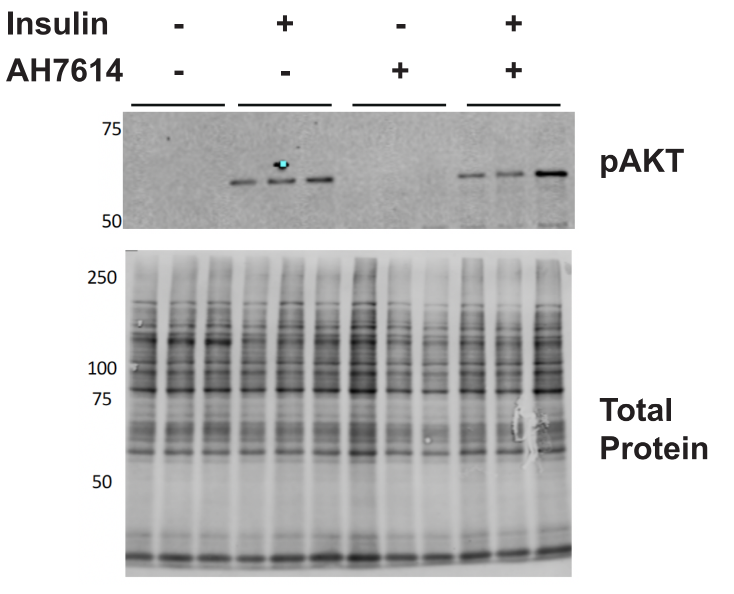
